# A performance evaluation of neural network features and functions settings on the model accuracy

**DOI:** 10.1101/2022.11.28.518263

**Authors:** Michal Bozděch

## Abstract

Not only in sports is a neural network the most used type of artificial intelligence. With software development, anyone can create a neural network model, but little is known about how to prepare the data and how to set up the model algorithms to their maximum performance. For these reasons, this study aims to determine whether features or function settings have a greater effect on model accuracy. An initial feature dataset (*n* = 18882) was obtained from publicly available sources. Each of the six different feature settings consisted of 96 models. A total of 384 models were created, in which their testing accuracy and the percentage difference between the training and testing phases were further analyzed. No statistically significant differences were found between the accuracy of the function’s settings, but statistically significant differences were confirmed between the feature settings. The study found that feature settings, especially the reduction of the number of outputs, are a more important factor in increasing the model accuracy, than function settings. Although the literature focuses more on the function setting and sets feature setting is taken rather as a type of how to improve the model.

## Introduction

With the growing general public’s interest in sports and technological development, a huge amount of structured and unstructured data is emerging that can be used for predicting future events [1]. Although, artificial intelligence (AI) has been used for over 20 years is still considered a relatively new (but rapidly evolving) technology capable of simulating human intelligence [2]. Machine learning (ML) is an integral part of AI and refers to the automated detection of patterns in datasets [3]. ML was first characterized as a field of study, that allows computers to learn without being explicitly programmed [4]. The newer definition characterizes ML as a computer programming process to optimize performance criteria using (mostly) data [5]. An increased interest in predicting sports outcomes started around 2010 [1]. ML is typically classified as supervised learning, unsupervised learning, and reinforcement learning. It is also possible to encounter an additional division into Semi-supervised learning, Transduction, and Learning to learn algorithms and more [5–7]. This study is a typical Supervised learning type of ML because the model learns from a labeled dataset. More precisely classification models, because outputs are discrete, instead of continuous outputs as in the case of regression models [6,8]. The correct use of model selection, evaluation, and algorithm selection techniques of machine learning is vital in the commercial sector as well as in academic research [9]. In sports, ML algorithms can help with predicting team’s and individual performance (in real-time but also as long-term forecasts), making sport-betting decisions, predicting career trajectory, identifying talented athletes and tactical patterns, preventing possible player injury, and so on [1,10–12].

Koseler and Stephan [13] found that Support Vector Machine and *k*-nearest neighbor’s were the most used ML algorithms used in baseball analytics. They also concluded that artificial neural networks (ANN) will become the most used model (in baseball). Which was confirmed (not only in baseball) by Horvat and Job [1] and Bunker and Susnjak [14] who also concluded that ANN does not necessarily perform significantly better than other ML models. However, it is possible to claim that each predictive model has its advantages and disadvantages and that it is not appropriate to compare the results of different studies, as they were obtained from different datasets [1,15].

An important preliminary step, that significantly contributes to the model accuracy is feature selection and extraction. This can be described as a process of identifying, adding, and removing irrelevant and redundant features and thus making the model faster, more efficient, and more accurate [16,17]. As already mentioned, adding features to the model (such as a combination of two other features; goals per number of played games) is another technique for maximizing the relevance of the model [18]. However, it is important to know that various ML algorithms with a higher number of features can suffer from overfitting and low accuracy [19]. Although there are various methods for reducing feature dimensions, it is difficult to conclude which method provides the best results [1]. Similarly, Raschka [9] further states that machine learning involves a lot of experimentation to gradually fine-tune the model’s accuracy. In the best situation, it does not experiment with only one algorithm (the best/logical for the given situation), but several, which are compared with each other. This is possible because they are data from the same dataset. In the past, only wealthier sports clubs could afford to pay a specialist (employees or services of another company) for data analysis and so they were able to (successfully) apply the results in practice. With available literature, the development of software with a simple and intuitive interface (such as SPSS, MATLAB, Statistica, etc.) the situation changes, and every sports manager (coach, stakeholder, data analyst, or sports enthusiast) can use AI to their advantage, without needing to know a suitable programming language and paying for data analyst services. Although it is possible to find exceptions [9,20–24] most studies do not present a comprehensive description of prediction (via accuracy) results from the comparison (as this is the know-how of the ML architect) using the initial feature set and feature selection, as adding and removing irrelevant and redundant features or compared accuracy results to different model settings, according to the number of hidden layers, types of activation function, output layers, types training and optimization algorithms. For these reasons, this study aimed to demonstrate the application of various methods to increase the accuracy of the predicted results of the data without the use of a specialized computer/programming language (coding). This procedure can be used by students, athletes’ managers’ and coaches of all clubs, not only specialized staff, which most clubs cannot afford for financial reasons.

## Methods

The dataset contained 18882 data points from a total of 1929 players. Obtained from the publicly available and validated (from the sports association) website (www.fortuna.cz). Inclusion criteria: seasons 2014/2015–2018/2019, the overall statistic of soccer players (of 16 teams) for the Czech highest league; only forward, defender, and midfielder playing positions. Excluded variables were playing positions goalkeeper, as their stats differed significantly from other player positions (e. g. height, number of goals, yellow and red cards received), variable nationality, and number of clean sheets (the variable related only to goalkeepers). Missing data were omitted. Due to the change in the rules and the objectives of this study, only the above seasons were analyzed.

Dependent variables (for the 1^st^ and 2^nd^ phases of this study) were nominal (player positions), ordinal (number of games played, numbers of goals, yellow and red cards received) and scale (age, height, weight) and represent the values at the end of the season. Five consecutive seasons have been analyzed separately (not cumulatively) as it provides greater accuracy [1].

Final team/players ranking at the end of the season (ordinal scale; 1st to 16th rank for 1^st^ to 4^th^ phases; by quarters for 5^th^ phase; 1^st^ rank and the Others ranks was used for 6^th^ phase) was the only output variable.

The default settings of the Artificial Neural Network (ANN) if there was only 1 hidden layer or a Deep Neural Network (DNN), if there were 2 hidden layers, were always set as Multilayer Perceptron (suitable for solving non-linear problems); Standardized Rescaling of Covariates; 70% of partitions divided into training and 30% to testing the model; the maximum number of epochs were 10,000. Where output (dependent variable) was the players/teams’ rank at the end of the season (discrete or ordinal variable) and input where decimal age calculates on the last day of the season (continuous variable), player’s height (continuous variable) and weight (continuous variable), number of played games (discrete variable), number of goals (discrete variable), number of yellow (discrete variable) and red cards (discrete variable) received. Additionally (in the 3^rd^ phase) the ratios were calculated from the available variables’ Goal/games, Goal/yellow cards, Goal/red cards, and Age/games.

The research contained 6 individual computed and analyzed phases of ML modeling using ANN/DNN, in which inputs and outputs features were gradually, systematically, and purposefully manipulated. (1) Only raw data (Raw_data) - without any modifications - was used to create the ANN and DNN models. (2) ANN and DNN models without any Outliers values were calculated. (3) The newly calculated variables were added to the model (Added_features). (4) Next, variables with the least effect on model accuracy were removed from the model (Removed_features). The last two phases did not deal with setting inputs, but outputs. (5) First in the form of a reduction from 16 possible outputs to 4, according to the quarter of the final order (Output_mod_1). (6) Last by reducing the number of outputs to 2 (1^st^ and the other ranks). This phase (outputs_mod_2) contained 16 cases/models where analysis could not be performed because one or more cases in the testing sample had dependent variable values that do not occur in the training sample. Although, making accurate ML algorithms do not have a one-size-fits-all approach this is the recommended procedure for refining model accuracy [1,25].

Each of the six individual phases contained the same and predetermined order set of the individual ANN/DNN model’s function. More precisely it was 2 hidden layers, 2 activation functions (Hyberbolic tangent, Sigmoid), 4 output layers (Identity, Softmax, Hyberbolic tangent, Sigmoid), 3 types of training (Batch, Online, Minibatch), and 2 optimization algorithms (Scaled conjugate gradient, Gradient descent). If algorithms that cannot be combined are not counted. There was a total of 64 different models in each phase.

Training and testing accuracy (%) were obtained for all calculated models (*n* = 384). Along with the results of the percentage difference (%diff) between training (V1) and testing (V2) accuracy.

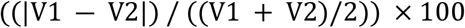

The most accurate models (from the testing accuracy) were more thoroughly analyzed using The Receiver Operator Characteristic (ROC), Area under the ROC Curve (AUC), and according to their percentage of importance of individual variables.

The assumption for the parametric analysis of variance test was violated according to the results of Levene’s test. Kruskal-Wallis H test was used to compare model accuracy results between different function settings (up to 96 individual models) and feature settings (1st to 6th phases of research). The level of statistical significance was set at *p* < 0.05. But was also adjusted for multiple comparisons using Bonferroni correction. IBM SPSS Statistics software version 28.0.0 for Windows (IBM SPSS Inc. Chicago, IL, USA) was used for the statistical analyses and to calculate the models. Except for the basic use of syntax, no coding was used during this study.

## Results and Discussion

Since we have 2 alternatives for hidden layers; 2 for the activation function; 4 for the output layer; 3 for a type of training; 2 for the optimization algorithm. There was a total of 96 different ways to modify a given model. The testing accuracy of values from the individual models, depending on the individual changes, are presented in Fig 1 and Fig 2 shows the percentage difference between the training and testing phases. Table 1 contains only the models with the lowest and the highest possible accuracy, gradually according to the recommended procedure for working with data when creating an ANN/DNN model: (1) raw data only; (2) after removing outliers from continuous variables; (3) adding possible relevant features; (4) removing redundant and irrelevant features (based on previous importance results); (5) modification of the number of outputs (based on the quarter of final rank at the end of the season). Results of the last phase (Output mod_2) are not listed in Fig 1 and Table 1 but are more thoroughly analyzed separately in text and Table 2.

**Fig 1.**
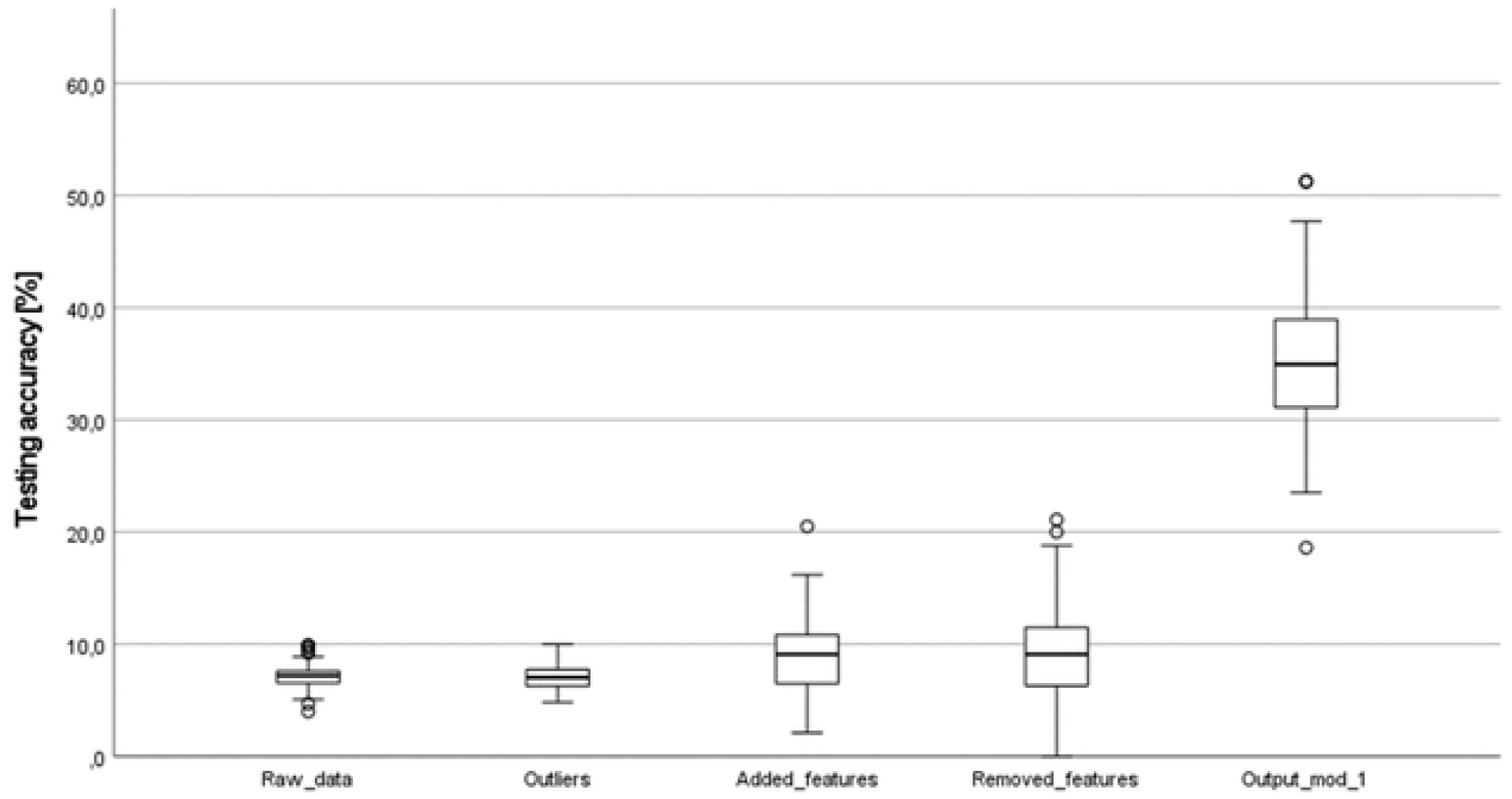
An overview of accuracy results from model testing in individual phases.

**Fig 2.**
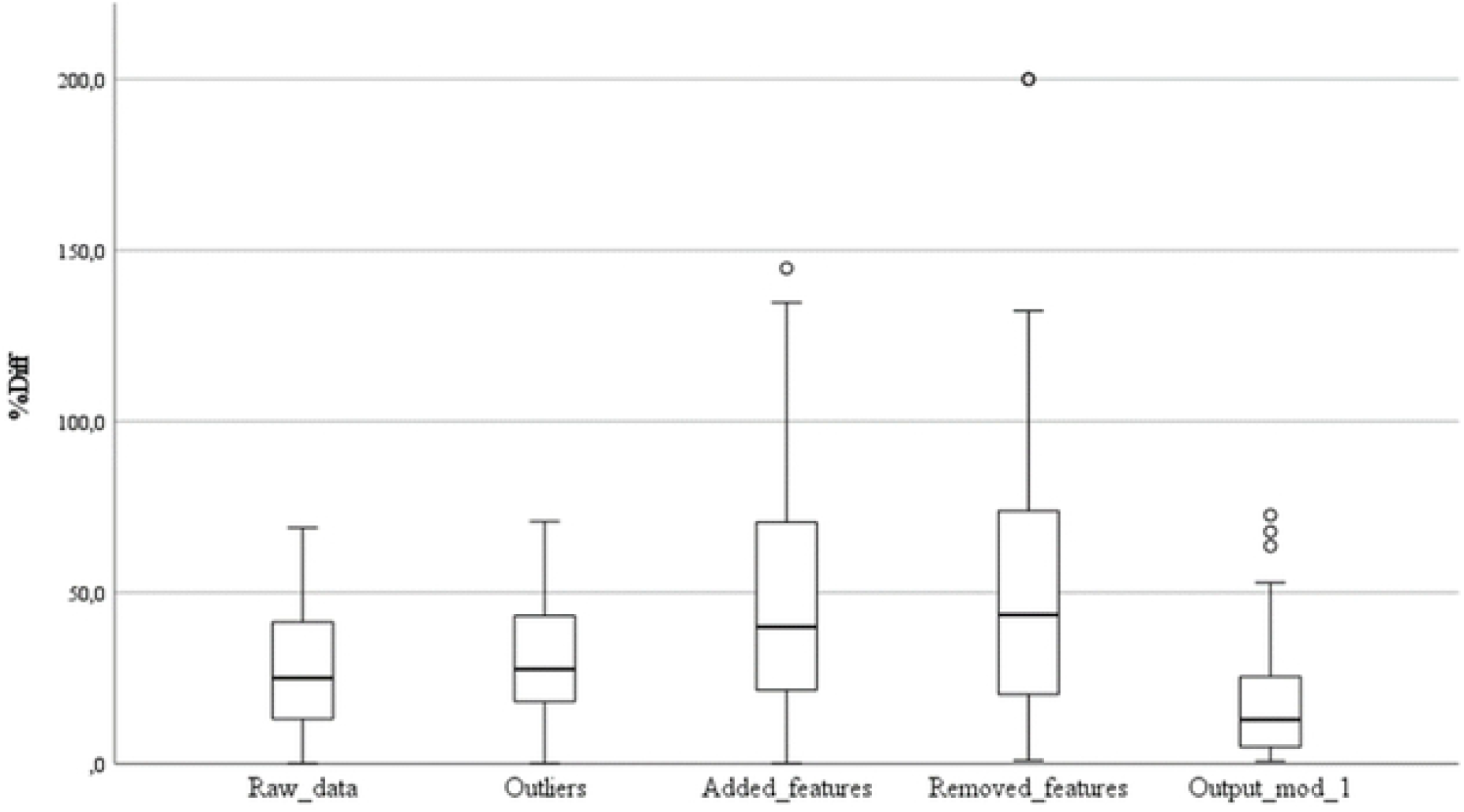
The percentage difference between the results of training and testing accuracy.

**Table 1.** An overview of the most accurate models in the four individual phases of strengthening.

| Model | Architecture |  |  | Training |  | Classification [%] |  |
| --- | --- | --- | --- | --- | --- | --- | --- |
|  | Hidden Layer | Activation Function | Output Layer | Type of Training | Optimization algorithm | Training | Testing |
| (1) Raw LA | 2 | Tanh | Tanh | Batch | GD | 8.2 | 4.0 |
| (1) Raw MA | 1 | Tanh | Identity | Minibatch | GD | 12.1 | 9.9* |
| (2) Outliers LA | 2 | Tanh | Identity | Online | GD | 8.4 | 4.8 |
| (2) Outliers MA | 1 | Tanh | Identity | Online | GD | 10.9 | 10 |
| (3) Add features LA | 1 | Tanh | Tanh | Online | GD | 13.1 | 2.1 |
| (3) Add features MA | 2 | Tanh | Identity | Minibatch | GD | 8.8 | 20.5 |
| (4) Remo. features LA | 1 | Sigmoid | Sigmoid | Batch | GD | 11.6 | 0.0* |
| (4) Remo. features MA | 2 | Sigmoid | Identity | Batch | SCG | 7.8 | 21.1 |
| (5) Output mod 1 LA | 2 | Tanh | Softmax | Batch | SCG | 39.8 | 18.6 |

LA = The least accurate model; MA = the most accurate model; Tanh = Hyperbolic tangent; SCG = Scaled conjugate gradient; GD = Gradient descent; \* = in case of duplicate results, larger or smaller values from the model training decided.

**Table 2.** Overview of the model settings with 100% accuracy and 1.00 AUC.

| Model | Architecture |  |  | Training |  |
| --- | --- | --- | --- | --- | --- |
|  | Hidden Layer | Activation Function | Output Layer | Type of Training | Optimization algorithm |
| [1] | 1 | Hyberbolic tangent | Softmax | Batch | Scaled conjugate gradient |
| [2] | 1 | Hyberbolic tangent | Softmax | Batch | Gradient descent |
| [3] | 1 | Hyberbolic tangent | Sigmoid | Batch | Scaled conjugate gradient |
| [4] | 1 | Sigmoid | Softmax | Batch | Gradient descent |
| [5] | 1 | Sigmoid | Sigmoid | Online | Gradient descent |
| [6] | 1 | Sigmoid | Sigmoid | Minibatch | Gradient descent |
| [7] | 2 | Hyberbolic tangent | Softmax | Batch | Scaled conjugate gradient |
| [8] | 2 | Hyberbolic tangent | Softmax | Online | Gradient descent |
| [9] | 2 | Hyberbolic tangent | Softmax | Minibatch | Gradient descent |
| [10] | 2 | Hyberbolic tangent | Sigmoid | Batch | Scaled conjugate gradient |
| [11] | 2 | Hyberbolic tangent | Sigmoid | Batch | Gradient descent |
| [12] | 2 | Hyberbolic tangent | Sigmoid | Minibatch | Gradient descent |
| [13] | 2 | Sigmoid | Softmax | Online | Gradient descent |
| [14] | 2 | Sigmoid | Softmax | Minibatch | Gradient descent |

The box plot in Fig 1 shows different dispersion of testing accuracy according to various features setting (phases). This could be described as the first phase containing the lowest prediction accuracy, which tended to increase with each sequential phase of modeling the ML algorithm. However, with increasing model accuracy the heterogeneity of the results also increased.

From Fig 1 can be seen that with each subsequent phase, which was done according to the recommendations of the literature [1,25], the accuracy of the model prediction increased. Which could be described as the correct applied setting of the model. However, because the percentage of differences between the training and testing phases gradually increased, until the output setting (see Fig 2), a model can be also described as overfitting [1,19]. Which was reduced by the 5^th^ and 6^th^ phases, more precisely, by reducing the number of outputs. Therefore, the reduction of the possible outputs could be described as a suitable technique not only to increase the accuracy of the model prediction but also to reduce the chances of model overfitting.

The increasing accuracy of the model with the increasing number of features contradicts Horvat and Job [1] conclusions, who found that with an increasing number of features (and season) the outcome prediction decreased. This opposite conclusion may be because the model from this study has not yet reached its greatest accuracy. Further in Fig 1 and 2 can be seen, that after removing the least important features from the model, neither the accuracy nor the difference changed significantly. According to the results of the descriptive analysis, it can be argued that the accuracy (of the testing phase) is similar between raw data (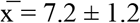; skewness = 0.1; kurtosis = 1.0) and data without outliers (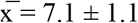; skewness = 0.2; kurtosis = −0.2). It increases slightly when features are added (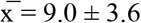; skewness = 0.5; kurtosis = 0.7), but does not change much after removing excess features (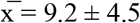; skewness = 0.4; kurtosis = 0.2). A big increase in accuracy came after the first reduction of outputs (from 16 to 4 possible outputs; Output_mod_1) variables (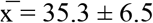; skewness = 0.1; kurtosis = 0.3). It is not necessary to characterize descriptive statistics of the last phases of this study (Output_mod_2), as all calculated results of model testing reached 100% accuracy.

From the above, it is possible to confirm Horvat and Job [1] conclusions that adding a feature (e.g. combining initial features) contributes to greater accuracy of the model. But a much more significant increase in model accuracy is achieved by reducing the number of possible outputs. This is confirmed by the Bunker and Susnjak [14] study, in which the authors state that sports with a larger number of possible outcomes provide generally a lower model accuracy. Which was most evident in the 6th phase of the study. Where the dependent variable was adjusted from four to two possible outputs. This caused 41 models to reach 100% accuracy (in testing). Weight (*n* = 15; 36.59%) was the variable with the highest number of occurrences of 100% Normalized importance. Then Height (*n* = 7; 17.07%), Age (*n* = 6; 14.63%), Goal/red (*n* = 4; 9.76%), Games played (*n* = 3; 7.32%), Age/games (*n* = 3; 7.32%), Goal/yellow (*n* = 2; 4.88%), number of goals (*n* = 1; 2.44%), Goal/games (*n* = 0; 0.00%), respectively. In the case of another phase (Removed_features_2) of model accuracy refinement, the last 3 variables would be excluded from modeling. The number of goals has been only found once as the feature with the highest normalized importance because in soccer (compared to other sports) fewer goals/points are scored or because all player positions have been added to the algorithm. It may be appropriate in future studies to calculate a separate algorithm for each player’s position and compare the occurrence of their Normalized importance.

After the fine-tuning of model accuracy (also called classification accuracy), it is typical that another type of performance metric optimization is performed, such as the Area Under the Curve (AUC) curve and the Receiver Operating Characteristic (ROC) curve [9]. They characterize how much the model can distinguish between outcomes (classes). Therefore these 41 most accurate models were further evaluated using ROC Curve and AUC, which ranged from 0.125 to 1.000 (AUC). The following Table 2 provides an overview of models (*n* = 14) that achieved 100% accuracy and AUC was equal to 1.00.

It seems that the best models are composed of two hidden layers and Hyberbolic tangent as the activation function. For the output layer, it does not matter whether you use the Softmax or Sigmoid activation function. For the type of training and optimization algorithm, the most appropriate solution was not found. Therefore, it maybe depends more on the data and the situation. Interestingly, Table 2 does not contain an Identity and Hyperbolic tangent output layer.

Activation functions transform a neuron’s activation level into an output signal. Several activation functions can significantly affect model performance/accuracy [24]. Studies focusing on the model accuracy according to activation function indicate that although the Hyperbolic Tangent Function is similar to the Sigmoid function, Sigmoid functions are the most commonly used and that different settings of the activation function improve ANN accuracy [24,26,27]. For DNN, Hyperbolic tangents as activation functions for both neurons (or nodes) prove better accuracy, which is confirmed by the results of this study and Karlik and Olgac [25].

The fact, that there is not one but exactly 14 models with 100% accuracy and 1.00 AUC was very unexpected. For comparison, the authors Hucaljuk and Rakipović [29] used several ML algorithms to predict outcomes in Champions’ League group stage games and achieved maximum accuracy of 68.8%. Similar accuracy was described by Horvat and Job [1] in their extensive review. Where the average maximum accuracy of predicting soccer matches outcome was 67.42%, without outliers (72.43% with 2 outliers), with a 54.55–93.00% range. In comparison, soccer’s prediction of match outcomes accuracies is lower compared to other sports. Because it is easier to predict the results of individuals than teams [1]. Where outcomes are affected by multiple variables, that are difficult or at all possible to measure. Another reason why the accuracy of the model was higher than other studies could be based on the fact, that a classification type (not regression type) model was used to predict the outcome [29–31]. It is important to note that it is not appropriate to compare the maximum accuracy of our model with others and it only serves as a supplement to the general idea of the issue. As their model was focused on predicting match (not season) outcomes, they come from different datasets or it is a model based on data from different sports [1,14].

Fig 1 indicated significant differences between the individual (1^st^ to 6^th^) research phases, which was confirmed by the Kruskal-Wallis test (*p* < .001). After adjusting for multiple comparisons testing the post hoc test also found statistically significant differences in 8/15 cases (53.3%). More precisely between Output_mod_1 and every other phase (except Output_mod_2) and Output_mod_2 and every other phase (except Output_mod_1). No statistically significant differences were found between the testing accuracy and the type of model setting (*p* > .05) in any of the phases. These results suggest that since there were no statistical differences in accuracy between the settings of the individual algorithms (in each phase), feature setting (especially the output variables) is more important for greater model accuracy. Which can also speed up the performance of the algorithm. But that was not the subject of this research.

This study does not replace the understanding of when to use individual settings (algorithms, functions). It only quantifies different features and function setting on model accuracy using one data set.

As mentioned above the neural network is the most widely used ML model. In practice, a neural network (ANN/DNN) may not be the most appropriate method for this case. Because the dataset consists of end-of-season data only (when the results are already known). Therefore, it would be appropriate to try other methods and procedures for predicting the final ranking of the team. Therefore, methods such as Forecasting (predicting changes over time) and Bayes Theorem, when the data set would be updated weekly, should provide more relevant predictions.

The inability to compare the properties and accuracy of two different models (since they do not come from the same dataset) significantly complicates the work of researchers and data engineers. Therefore, it would be appropriate to invent or derive some methods, approaches, or strategies for this type of analysis. For example, based on the effect size testing.

## Conclusions

Due to the huge popularity of artificial intelligence and machine learning, the number of statistical software, where it is possible to create an ML model without knowing the coding language, is growing. For these reasons, this study focused on quantifying the effect of feature and function settings on the accuracy of the classification type model. Study results suggest that, since there were no differences in model accuracy between the settings of the individual algorithms (in every individual phase), In contrast to the feature settings where statistically significant differences were found. We can state that feature settings are more important for better model accuracy. In particular, a lower number of output variables has the best positive effect on model accuracy. Therefore, it would be appropriate to prepare the model features and consider with an expert on a given sports issue what the model should predict and whether the results of the model are applicable in practice. From a total of 41 different algorithms which reached 100% model accuracy weight, height and age were the most often feature with normalized importance occurrences. This suggests that not the number of goals (or other derivatives arising from this variable), but these variables are better for predicting whether a team will win the league.

## Supporting information

**S1 Table. Overview of the most accurate models in the four individual phases of strengthening.**

LA = The least accurate model; MA = the most accurate model; Tanh = Hyperbolic tangent; SCG = Scaled conjugate gradient; GD = Gradient descent; * = in case of duplicate results, larger or smaller values from the model training decided.

## References

1. Horvat T, Job J. The use of machine learning in sport outcome prediction: A review. Wiley Interdiscip Rev Data Min Knowl Discov. 2020;10. doi:10.1002/widm.1380

2. Glikson E, Woolley AW. Human Trust in Artificial Intelligence: Review of Empirical Research. Academy of Management Annals. 2020;14: 627–660. doi:10.5465/annals.2018.0057

3. Woschank M, Rauch E, Zsifkovits H. A review of further directions for artificial intelligence, machine learning, and deep learning in smart logistics. Sustainability (Switzerland). 2020;12. doi:10.3390/su12093760

4. Samuel AL. Some Studies in Machine Learning Using the Game of Checkers. IBM Journal. 1959;3: 535–554.

5. Alpaydim E. Introduction to Machine Learning. The MIT Press; 2010.

6. Nasteski V. An overview of the supervised machine learning methods. Horizons. 2017;4: 51–62. doi:10.20544/HORIZONS.B.04.1.17.P05

7. Kotsiantis SB. Supervised Machine Learning: A Review of Classification Techniques. Informatica. 2007;31: 249–268.

8. Gianey HK, Choudhary R. Comprehensive Review On Supervised Machine Learning Algorithms. Proceedings - 2017 International Conference on Machine Learning and Data Science, MLDS 2017. Institute of Electrical and Electronics Engineers Inc.; 2018. pp. 38–43. doi:10.1109/MLDS.2017.11

9. Raschka S. Model Evaluation, Model Selection, and Algorithm Selection in Machine Learning. 2018. Available: http://arxiv.org/abs/1811.12808

10. Barron D, Ball G, Robins M, Sunderland C. Artificial neural networks and player recruitment in professional soccer. PLoS One. 2018;13. doi:10.1371/journal.pone.0205818

11. Muazu Musa R, Abdul Majeed APP, Taha Z, Abdullah MR, Husin Musawi Maliki AB, Azura Kosni N. The application of Artificial Neural Network and k-Nearest Neighbour classification models in the scouting of high-performance archers from a selected fitness and motor skill performance parameters. Sci Sports. 2019;34: e241–e249. doi:10.1016/j.scispo.2019.02.006

12. Pfeiffer M, Hohmann A. Applications of neural networks in training science. Hum Mov Sci. 2012;31: 344–359. doi:10.1016/j.humov.2010.11.004

13. Koseler K, Stephan M. Machine Learning Applications in Baseball: A Systematic Literature Review. Applied Artificial Intelligence. 2017;31: 745–763. doi:10.1080/08839514.2018.1442991

14. Bunker R, Susnjak T. The Application of Machine Learning Techniques for Predicting Results in Team Sport: A Review. 2019. doi:10.1613/jair.1.13509

15. Haghighat M, Rastegari H, Nourafza N. A Review of Data Mining Techniques for Result Prediction in Sports. Advances in Computer Science: an International Journal. 2014;2: 7–12. Available: https://www.researchgate.net/publication/262560138

16. Han J, Kamber M, Pei J. Data Mining: Concepts and Concepts and Techniques. 3rd ed. Waltham: Elsevier; 2012.

17. Yu L, Liu H. Efficient feature selection via analysis of relevance and redundancy. The Journal of Machine Learning Research. 2004;5: 1205–1224.

18. Jović A, Brkić K, Bogunović N. A review of feature selection methods with applications. Proceedings of 38th international convention on information and communication technology, electronics and microelectronics (MIPRO). Opatija; 2015. doi:doi: 10.1109/MIPRO.2015.7160458.

19. Lin J, Short L, Sundaresan V. Predicting National Basketball Association Winners. 2014.

20. Horvat T, Job J, Medved V. Prediction of euroleague games based on supervised classification algorithm k-nearest neighbours. Proceedings of the 6th International Congress on Sport Sciences Research and Technology Support (icSPORTS 2018). SciTePress; 2018. pp. 203–207. doi:10.5220/0006893502030207

21. Ping-Feng P, Lan-Hung C, Kuo-Ping L. Analyzing basketball games by a support vector machines with decision tree model. Neural Comput Appl. 2017;28: 4159–4167. doi:10.1007/s00521-016-2321-9

22. Trawiński K. A fuzzy classification system for prediction of the results of the basketball games. 2010 IEEE World Congress on Computational Intelligence, WCCI 2010. 2010. doi:10.1109/FUZZY.2010.5584399

23. Loeffelholz B, Bednar E, Bauer KW. Predicting NBA Games Using Neural Networks. J Quant Anal Sports. 2009;5. doi:10.2202/1559-0410.1156

24. Karlik B, Olgac AV. Performance Analysis of Various Activation Functions in Generalized MLPArchitectures of Neural Networks. nIternational Journal of Artificial Intelligence and Expert Systems. 2011;111. Available: https://www.researchgate.net/publication/228813985

25. MathWorks. Introduction to Machine Learning, Part 4: Getting Started with Machine Learning. 4 Mar 2020 [cited 11 Jul 2022]. Available: https://www.mathworks.com/videos/introduction-to-machine-learning-part-4-getting-started-with-machine-learning-1542879650900.html

26. Widrow B, Lehr MA. 30 Years of Adaptive Neural Networks: Perceptron, Madaline, and Backpropagation. Proceedings of the IEEE. 1990;78: 1415–1442. doi:10.1109/5.58323

27. Buhmann MD. Radial basis functions: theory and implementations. Cambridge university press; 2003.

28. Hucaljuk J, Rakipović A. Predicting football scores using machine learning techniques. Proceedings of the 34th International Convention MIPRO. 2011. Available: https://ieeexplore.ieee.org/stamp/stamp.jsp?tp=&arnumber=5967321&isnumber=5967009

29. Delen D, Cogdell D, Kasap N. A comparative analysis of data mining methods in predicting NCAA bowl outcomes. Int J Forecast. 2012;28: 543–552. doi:10.1016/j.ijforecast.2011.05.002

30. Elfrink T. Predicting the outcomes of MLB games with a machine learning approach. Vrije universiteit. 2018.

31. Soto Valero C. Predicting win-loss outcomes in MLB regular season games-a comparative study using data mining methods. Int J Comput Sci Sport. 2016;15: 91–112. doi:10.1515/ijcss-2016-0007

